# A unified theory of distraction in human perceptual, cognitive and economic decision-making

**DOI:** 10.1101/160143

**Authors:** Vickie Li, Elizabeth Michael, Jan Balaguer, Santiago Herce Castañón, Christopher Summerfield

## Abstract

When making decisions, humans are often distracted by irrelevant information. Distraction has different impact on perceptual, cognitive and value-guided choices, giving rise to well-described behavioural phenomena such as the tilt illusion, conflict adaptation, or economic decoy effects. However, a single, unified model that can account for all these phenomena has yet to emerge. Here, we offer one such account, based on adaptive gain control, and additionally show that it successfully predicts a range of counterintuitive new behavioural phenomena on variants of a classic cognitive paradigm, the Eriksen flanker task. We also report that BOLD signals in a dorsal network prominently including the anterior cingulate cortex (dACC), index a gain-modulated decision variable predicted by the model. This work unifies the study of distraction across perceptual, cognitive and economic domains.

## Introduction

Decisions about sensory signals, cognitive propositions, or economic prospects are often made in the context of competing or distracting information. Consider the following everyday situations: you are judging whether a painting hangs straight on the wall, but the nearby pictures are hung askew; you are waiting at a red stop signal, but the car in front decides to jump the light; you are contemplating the purchase of a new watch, but it is displayed next to a range of more elegant but unaffordable models. In each of these situations, the best decisions will be made by ignoring the distracting sensory signals (the competing picture frames, vehicles, or watches) and focussing exclusively on the choice-relevant information. This normative contention can be formalised in a variety of ways, for example via the notion that rational choices should be independent of irrelevant alternatives^1,2^ or that sensory signals should be weighted lawfully by their reliability and relevance to the choice at hand^3-6^.

Nevertheless, empirical observations suggest that humans decision-makers are unduly influenced by distracting information. Consider a generic problem in which a target stimulus *X^i^* and distracters *X^j^* occur at fixed locations *i* and *j*. In this general formulation, decision values *X* may be perceptual features (such as the tilt of a grating) or economic attributes (such as the quality of a consumer product) that are to be evaluated or categorised. Humans show systematic biases that reflect the influence of the distracters on decisions about the target. For example, vision scientists have long studied the “tilt illusion”, in which the reported orientation of *X^i^* (e.g. a central grating) is repulsed away from the mean tilt of *X^j^* (surrounding gratings with similar but nonidentical tilt; **Fig. 1a**)^7^. In cognitive psychology, the influence of distracter items is usually studied with a view to understanding the attentional or control mechanisms that allow information to be selected in the face of conflict. For example, in the classic Eriksen flanker task, observers classify a target stimulus (e.g. a central arrow) that is flanked by distracters (e.g. arrows pointing in compatible or incompatible directions)^8,9^. It is ubiquitously observed that incompatible flankers incur a cost, and compatible flankers a benefit, in response times (RTs) and accuracy relative to a neutral condition **(Fig. 1b)**. Finally, behavioural and neural economists have charted the irrational influence that a decoy alternative of value *X^j^* has on choices between two choice-relevant prospects *X^i+^* and *X^i-^*, where *X^i+^* > *X^i-10-13^*. A common finding is that rational choices (i.e. for *X^i+^* > *X^i-^*) initially decline as *X^j^* increases in value, but then increase sharply as *X^j^* comes to approximately match the other two items in value **(Fig. 1c)**.

**Figure 1.**
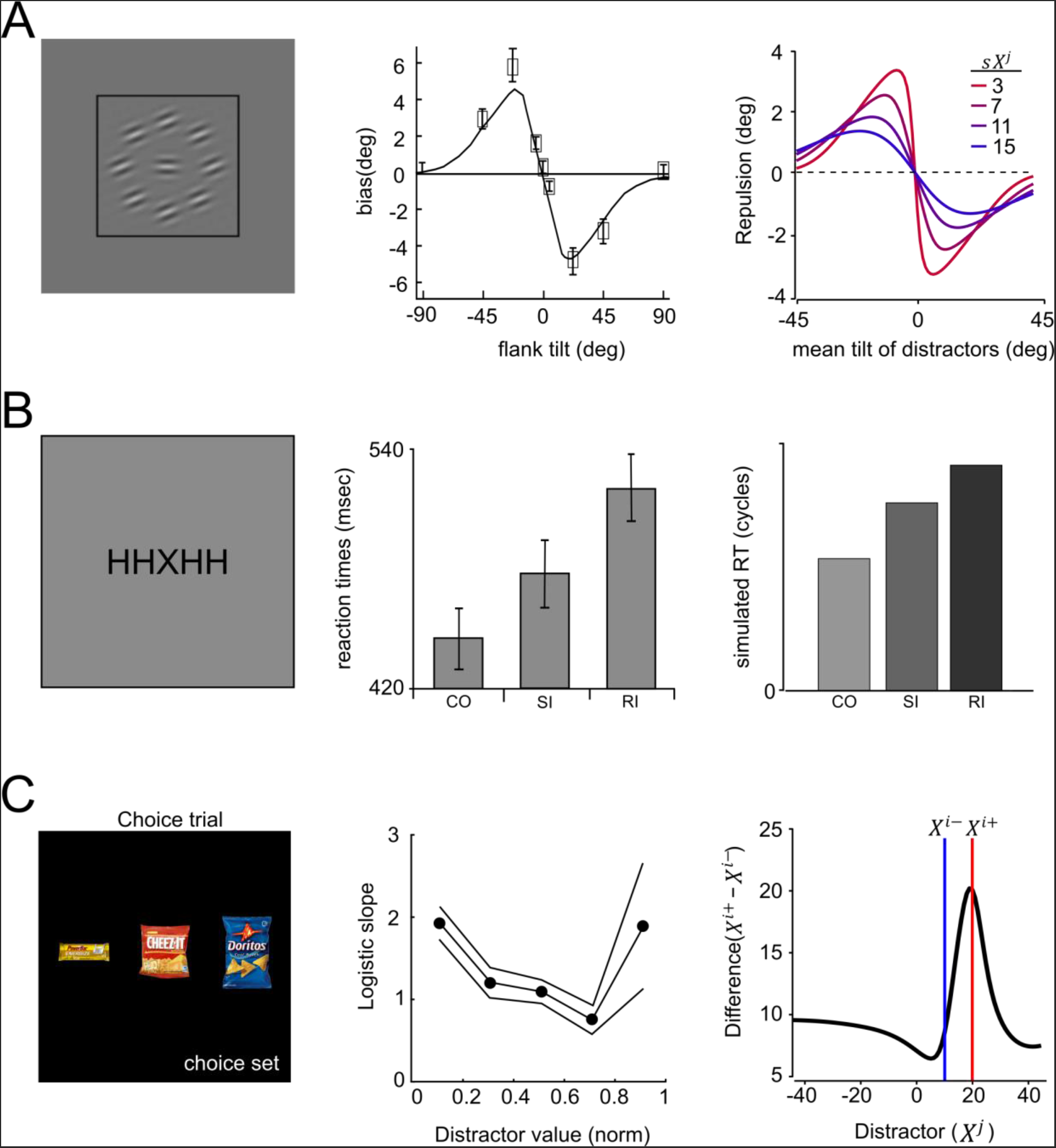
The effect distraction across perceptual, cognitive and economic domains. **(A)** Left panel: subjects were asked to discriminate the tilt (relative to horizontal) of a central Gabor surrounded by tilted distracters. Middle panel: participants are biased to report the target as more clockwise when the flankers were counterclockwise and vice versa (the ‘tilt illusion’, middle panel; images adapted from Solomon & Morgan, 2006, permission pending). Right panel: simulation of the adaptive gain model recreates the tilt illusion and further predicts that the magnitude of the bias is modulated by flanker variance; coloured lines reflect flanker standard deviation from low (red) to high (blue). **(B)** Left panel: In the Eriksen flanker task, participants respond with a key press to a central letter while ignoring the flankers. Middle panel: response times are the fastest on CO (‘congruent’) trials, then the SI (‘Stimulus Incongruent’) trials, and slowest on RI (‘Response incongruent’) trials; Middle panel; adapted from van Veen & Carter, 2002, permission pending). See methods for details of how CO, RI and SI trials were defined. Right panel: the adaptive gain model predicts the same pattern of reaction time across the three conditions. **(C)** Left panel: participants chose the most preferred of three food items. Middle panel: increasing the value of the least-preferred item reduces the choice efficiency (i.e. probability of choosing the highest-valued target) as the normalised distractor value increases, shown by logistic slope from fitting logistic choice functions on humans choice. Using the adaptive gain model, we simulated the subjective difference between two tilts (blue and red line) as a function of a third distracting tilt (x-axis). The subjective difference is first reduced and the increased in a qualitatively similar fashion.

In the fields of psychology, economics, and neuroscience, diverse theoretical proposals have been offered to explain the cost that distracters incur during decision-making. These include models that describe how control systems detect and resolve conflict among inputs^14,15^, accounts that emphasise inhibitory interactions among competing sensory neurons, or favour a normalisation of stimulus values by a local average or range^10,16-18^, and Bayesian accounts that model spatial uncertainty among targets and distracters^19-21^, or that assume strong priors on the compatibility of decision information^21^. These accounts disagree about the computational mechanisms involved, the neural processing stages at which the cost of distraction arises, and about the brain structures that are recruited to protect decisions against irrelevant information. For example, divisive normalisation mechanisms may occur in sensory neurons in visual cortex^16^, or amongst value representations in the orbitofrontal cortex^22^, whereas the control systems that detect and resolve conflict have been attributed to medial and lateral prefrontal structures^14^. As such, the field currently lacks a single, unified theory that can account for the effect of distraction on human decisions, or an integrated neural account of its implementation across perceptual, cognitive and economic domains.

The goal of the current paper is to offer such an account. We begin with a simple computational model that is motivated by past work showing that contextual signals determine the gain of processing of consistent (or “expected”) features during decision-making tasks^23^. Our model, which is described here at the level of neural population codes, proposes that contextual signals sharpen the tuning curves of neurons with compatible preference for decision-relevant features, and is motivated by a large literature emphasising the need for adaptive gain control in the service of efficient coding^24,25^. Using computational simulations, we first show that the model can recreate qualitatively two classic phenomena in very different domains – perceptual choice (the tilt illusion) and economic choice (decoy effects). Next, we turn our attention to a task that has been a mainstay of cognitive studies of distraction – the Eriksen Flanker task. We built novel variants of the task in which the statistics of the flankers, and the difference between target and the decision bound can vary across conditions. Our simulations show that the model predicts a range of striking, counterintuitive behavioural findings, including “reverse” compatibility effects (where fully-visible, compatible flankers actually hinder, rather than help, behavioural performance). Over 4 behavioural experiments involving human participants, we validate these predictions, using visual stimuli defined by both tilt and colour. Finally, we use functional brain imaging to show that the modulatory influence on decision signals predicted by the model correlates with BOLD signals the dorsal anterior cingulate cortex (dACC) and interconnected structures, where neural signals have variously been implicated in the context-sensitive encoding of action values^26^, and the expected value of cognitive control^27^. We show how our framework, which is not wholly inconsistent with either account, can bring together diverse views concerning the function of this controversial brain region^28^.

## Results

Our adaptive gain model is based on a framework that was previously developed to understand how humans performing perceptual decision-tasks adapt to the context provided by information that is proximal in space and time^23,29^. Inputs arrive at a population of *n* neurons each characterised by a Gaussian tuning curve centred on its preferred feature value *θ_k_*. Each neuron *k* responds to the target stimulus *X^i^* with rate *R_k_* = *f*(*X^i^*|*θ_k_, σ_k_*), where *f*(*X*|*θ,σ*) denotes the probability density function of the normal distribution with mean *θ* and variance *σ^2^*.

The estimated output of the neural population is then decoded into a subjective percept of the target 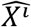 by weighting the population activity *R* by the corresponding feature values *θ*.

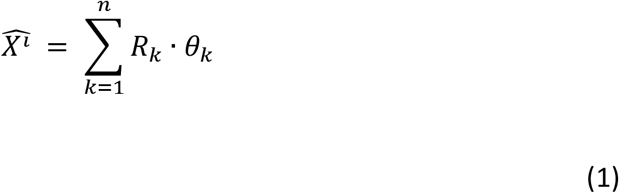

When the gain is uniformly spread across the feature space (i.e. the tuning widths *σ_k_* for all neurons are equal) this approach faithfully decodes each input to its original feature value. However, our model proposes that the context provided by the distracters modulates the sharpening of neuronal tuning, with a tuning width envelope that matches the inverse distribution of contextual features *X^j^* with mean *mX^j^* and standard deviation *sX^j^*.

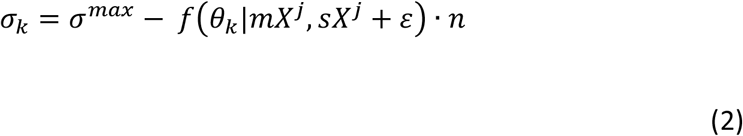

In other words, neurons with preferred orientation that matches *mX^j^* have the sharpest tuning curves, and these tuning curves are even sharper if the flanker variance (*sX^j^*) is low (see **Fig. 2a**). In equation (2), *σ^max^* denotes the maximum tuning width in the population, and *ε* is a noise parameter. In all model fitting procedures below, we allow *ε* to vary as a free parameter, recovering average values of ~5°, which concurs with previous accounts^30^.

**Figure 2.**
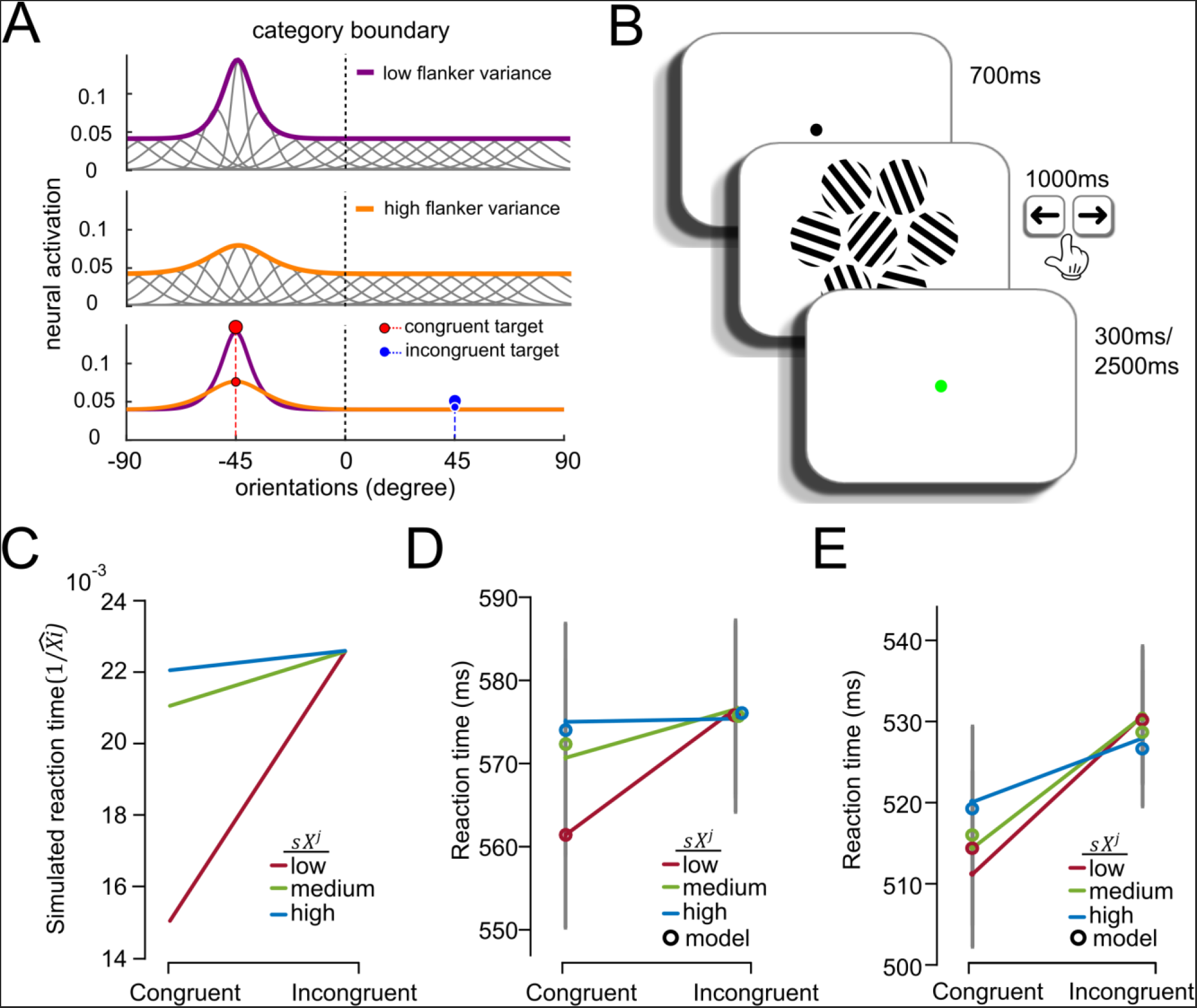
Adaptive gain model of the Eriksen flanker task. **(A)** Illustration of the adaptive gain model as applied to **Exp.1**. The model assumes that the population width tuning envelope is governed by the trial flanker mean (*mX^j^)* and trial flanker variability (*sX^j^*). Congruent targets (red circles) receive higher neural gain than incongruent targets (blue circles), leading to faster RTs. The model further predicts an interaction between congruency and flanker variability. Congruent targets receive high neural gain under low flanker variability trials (large circles) than high flanker variability trials (small circles) while incongruent targets under the two flanker variability trials received the same, low level of neural gain, meaning that flanker variability has no influence on RT for incongruent targets. **(B)** Experiment 1. Participant first saw a fixation dot, followed by an array that contained a central target and 6 surrounding flankers. They responded whether the central target was clockwise or counterclockwise to the vertical axis, receiving feedback after each response. **(C)** Simulation of RT 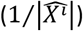 using the adaptive gain model for 3 levels of flanker variance (*sX^j^* in coloured lines) and congruency (x-axis). (**D - E**) Human reaction time pattern under three levels of flanker variance and congruency condition in **Exp. 1a** (Panel D) and **Exp.1b** (Panel E). Coloured circles represent the fitted mean reaction time from the adaptive gain model.

We first show how the model explains the both tilt illusion in perceptual choice tasks and decoy effects in economic choice tasks. In Fig. 1a, we plot the tilt bias over visual angle *θ* ∈ {-90,-89…90} predicted by the model as a difference of subjective and objective estimates of the target 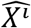 – *X^i^* for different values of *mX^j^* ∈ {-45,-44.45} and *sX^j^* ∈ {3, 7,11,15}. The model predicts that subjective estimates are repulsed away from the mean flanker value, as described in numerous previous studies^7,31^, and additionally that the strongest repulsion effect occurs when the flankers are consistent, i.e. are drawn from a distribution with low dispersion. This repulsion of the subjectively decoded values from their objective counterparts occurs when the target feature close to, but is not identical to, the mean of the distracters (i.e. the location of sharpest tuning), because the variable tuning profile induces a skew in the population activity over features *θ_k_* (see **Fig. S1** for a more detailed explanation). To model economic decoy effects, we envisage a choice between two prospects of value *X^i+^* and *X^i-^* (where *X^i+^* = 20 and *X^i-^* = 10 in arbitrary units) that is made in the context of distracters with average value *X^j^*. We plot the difference in their corresponding subjective estimates 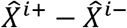 as a function of *mX^j^*, observing a pattern with a striking qualitative resemblance to that reported previously^10^ **(Fig. 1b)**. Again, the model’s ability to predict this counterintuitive pattern comes from the repulsive effect induced by differential tuning across feature space. The model predicts that as the value of the decoy *mX^j^* increases, repulsion is first strongest towards *X^i-^* (leading to a reduced preference for the best option *X^i+^*) but then, as the decoy approaches the two items in the choice set, repulsion is maximal for *X^i+^*, reversing this effect. In further simulations, we systematically varied both the distance between *X^i+^* and *X^i-^*, and *sX^j^*, we were able to capture the two-dimensional pattern of multi-alternative choice data described in a different study involving abstract shapes associated with different economic values^11^ (**Fig. S2**).

Next, we used our model to simulate performance on a new variant of the flanker task that involves categorising a central grating *X^i^* tilted at -45° from vertical, in the face of flanking gratings that are on average tilted in a compatible (*mX^j^* = -45°) or incompatible (*mX^j^* = +45°) fashion. **Fig. 2c** illustrates predictions from the adaptive gain model under two orthogonally varying factors: compatibility and flanker variability. Because flanker effects for fully-visible stimuli are strongest for response times, we plot the inverse model output 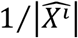 as a proxy for RT (this is equivalent to assuming a ballistic evidence accumulation process; see equation (1) and methods). As can be seen, the model predicts a compatibility effect: faster RTs for trials where the target and distracters were of congruent sign. However, it also makes a new, testable prediction: that as *sX^j^* (flanker variance) decreases, response times should be reduced on compatible trials but remain the same on incompatible trials **(Fig. 2c)**. This occurs because on compatible trials (flankers at – 45°), more gain is allocated to the target feature when the variance of the distribution of flanker orientations is lower. However, the gain allocated to incongruent targets (flankers at +45°) is negligibly different across different flanker variance levels since the neural gain they received are similar at the tail of the gain distribution, and so the model predicts that flanker variance should not affect performance on incongruent trials **(Fig. 2a)**. By contrast, classic models propose that response conflict varies with the amount of crosstalk interference among responses^14^. These models predict that heightened flanker variance should have equal impact on compatible and incompatible trials (see Methods for model details).

We tested this prediction in **Exp. 1** by asking healthy human participants to judge, relative to vertical, the tilt of a single target grating surrounded by 6 flanking distracter gratings **(Fig. 2b)**. The orientation of the target grating *X^i^* and the mean of the flankers *mX_j_* were set to ±45° and the standard deviation of the flankers *sX^j^* was varied at three levels: {0°, 15°, 30°} in **Exp. 1a** (n = 37) and {5°, 10°, 15°} in **Exp. 1b** (n = 36). In both experiments, we observed that flanker variance nodulated RTs on compatible trials (**Exp. 1a**: *F*_*1.91,68.6*_ = 9.041, *p* < 0.001; **Exp. 1b**: *F*_*1.91,68.86*_ = 20.16, *0* < 0.001) but not on incompatible trials (both p-values ≥ 0.09). This finding was qualified by an nteraction between compatibility and flanker variance (**Exp. 1a**: *F*_*1.83,65.79*_ = 12.14, *p* < 0.001; **Fig. 2d Exp. 1b**: *F*_*1.95,70.06*_ = 20.32, *p* < 0.001; **Fig. 2e**). Because previous work has shown that the ratio of compatible to incompatible flankers can modulate performance^32^, we repeated this analysis imited to those trials where all flankers fell on the compatible/incompatible side of the boundary, Finding a similar interaction for **Exp. 1a** (*F*_*1.83,66.04*_ = 12.59, *p* < 0.001) and **Exp. 1b** (*F*_*1.65,57.66*_ = 15.77, p < 0.001).

We fit our adaptive gain model to the data, and compared its predictions to those of a model oroposing that RT depends on response conflict alone. The fits for the gain model (coloured circles) are shown superimposed upon the human data in **Fig. 2c and 2d**. We compared the models head-to-head by computing mean-squared error (MSE) in RT across conditions on half of the data (even trials), after estimating parameters from an independent dataset (odd trials). Bayesian model selection showed that the adaptive gain model fit the human data more closely than the conflict model, with exceedance probabilities for the adaptive gain model of 0.95 in **Exp. 1a** and 0.72 in **Exp. 1b**. We also compared a version of the gain model in which the contextual modulation was driven by both target and distracters; this model yielded both qualitatively and quantitatively similar results to the original gain model (p > 0.2 for both **Exp. 1a** and **1b**).

Next, we moved beyond the simple case in which *X^i^* and *mX^j^* fell equidistant the category boundary to a more complex design where they could vary independently around vertical at ±15°, ±30°, ±45°), and *sX^j^* could once again vary at 3 levels {0°, 15°, 30°}. In this case, the model makes several predictions, some of them highly counterintuitive. Firstly, it predicts that there should be no main effect of congruence on RTs. In other words, the predicted inverse decision values 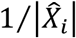 (or equivalently, unscaled RTs) are identical whether the target (*X^i^*) and flanker mean *mX^j^* are of the same and different sign. Secondly, the model predicts the existence of strong “reverse compatibility” effects under specific circumstances: there will be a disproportionate cost on congruent trials when the target *X^i^* is closer to the category boundary than the mean of the flankers *mX^j^*, e.g. when *X^i^* = 15 and *mX^j^* = 30 or *mX^j^* = 45 (**Fig. 3a** upper left corner of each plot), and that this effect should diminish with increasing flanker variance (panels). Finally, although the model indicates that RTs will be dominated by the distance between *X^i^* and the category boundary, it predicts that in the specific case where *X^i^* and *mX^j^* are both close to vertical, this cost will be strongly attenuated.

**Figure 3.**
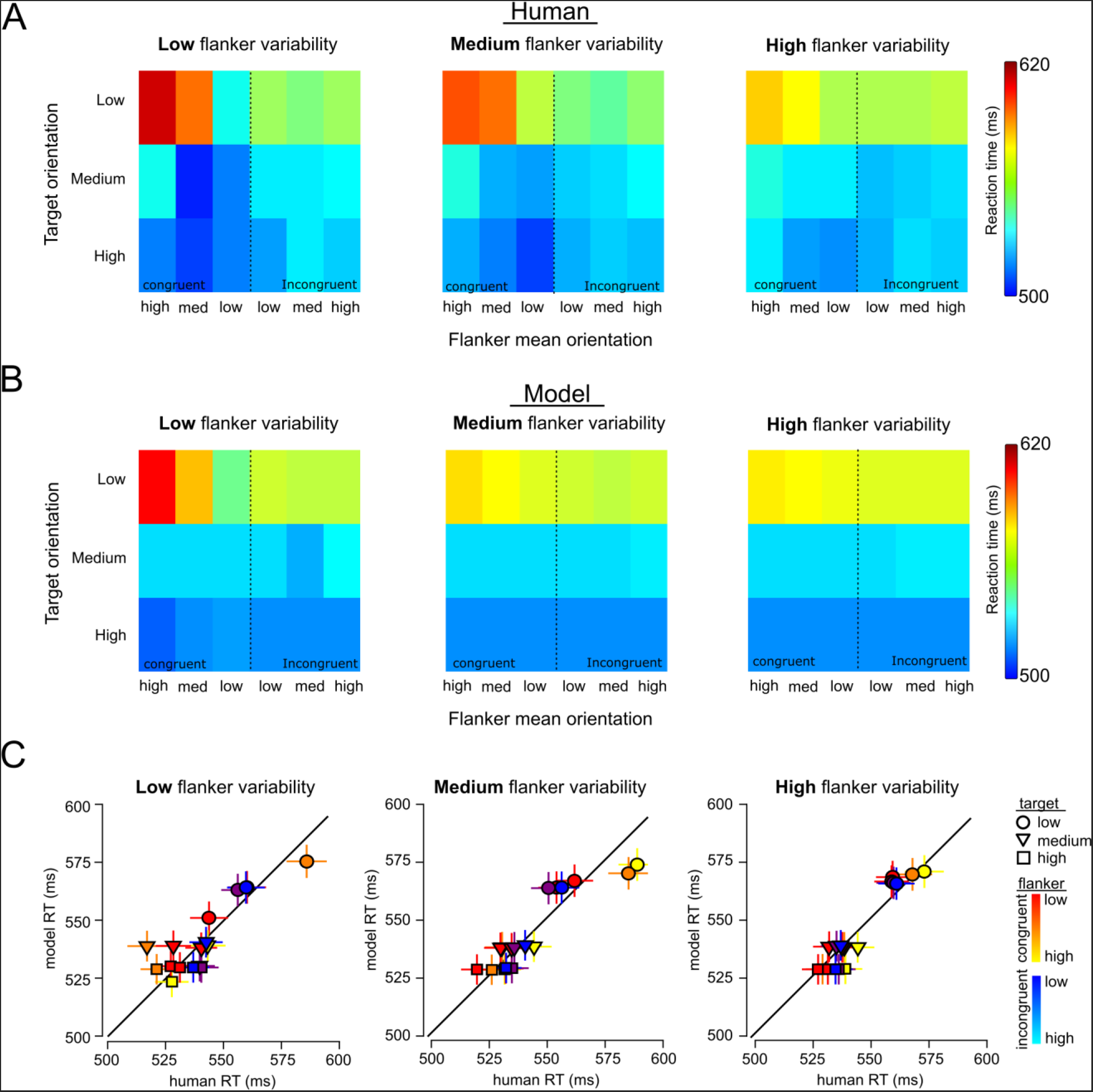
4 way interactions. **(A)** Surface plots showing the mean RT pattern in humans under different conditions (3 levels of target orientation x 3 levels of flanker mean orientation x congruency x 3 levels of flanker variability). Warmer colours correspond to longer reaction time. There is an overall cost when the target is close to the category boundary (top row of each surface plot). There is an additional cost when these targets were flanked by congruent flankers that are further from the boundary (top left corners of each subplot). This cost is radically reduced when the flankers are both congruent and close to the boundary (c.f. third most column, top row). The introduction of higher flanker variability reduces these additional costs in those conditions (overall faster RT across surface plots). **(B)** Fitted mean reaction time pattern from the adaptive gain model. **(C)** Mean (±SEM) RT in humans were cross-plotted against the fitted mean (±SEM) model RT for each condition. Warm colours (Red to yellow) correspond to levels of mean orientation from congruent flankers. Cold colours (Blue to cyan) correspond to levels mean orientation from incongruent flankers |*mX^j^*|. Different shapes correspond to 3 levels of target decision variable |*X^i^*|. Similar to upper plots, slower RT occurs specifically on trials where the target is close to the boundary and flankers are both congruent and have a mean orientation that is further from the category boundary.

We tested these predictions using the flanker paradigm on two new cohorts of participants, one of which (**Exp. 2a**, n = 28) performed the tilt categorisation task described above, except with the full 6 (target mean) x 6 (flanker mean) x 3 (flanker variance) design. Another (**Exp. 2b**, n = 30) performed a task with the same design that involved judging the colour of a central circle (red vs. blue) surrounded by distracting flankers that varied continuously in colour from red to blue. Results from the two experiments were qualitatively very similar (see **Fig. S3, S4** for separate data) and so after normalising the feature values (tilt, colour) to an equivalent scale in the range [-1,1] we collapsed over them for display purposes. All three model predictions were strongly present in the human RT data **(Fig. 3a-c)**. Firstly, splitting trials naively into compatible and incompatible, there was no overall significant difference in RT, either in the combined cohort (*t*_57_ = 1.14, *p* = 0.26) or separately for each experiment (**Exp. 2a**: *p* = 0.79; **Exp. 2b**: *p* = 0.17). Secondly, we see a strong cost on congruent trials when the flanker mean is further from the boundary than the target mean, as demonstrated by an *X^i^* × *mX^j^* interaction (*F*_*3.72,208.23*_ = 17.48, *p* < 0.001) that was much weaker on incongruent trials (*F*_*3.39,212.25*_ = 2.47, *p* = 0.049). This difference was qualified by a reliable three-way |*X^i^*| × |*mX^j^*| × *congruence* interaction on human RTs (*F*_*3.54,198.12*_ = 14.62, *p* < 0.001).

The analyses described thus far pertain to RT data. In a final analysis of **Exp.2** we examined choices, using a previously-described approach based on probit regression to assess the weight (or influence) that distracters wielded over choices, as a function of whether their tilt was inlying or outlying with respect to category boundary ^33,34^. We attempted to predict choices on each trial as follows:

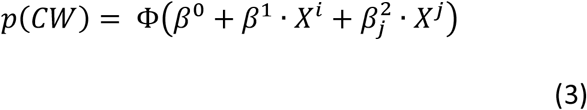

Where *X^j^* is tallied into 8 bins according to its signed feature value (e.g. from most counterclockwise to clockwise, or most red to blue). Plotting the coefficients 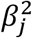 associated with each bin *j* for model choices (obtained via the fitting procedure above), the model counterintuitively predicts that inlying flankers (those whose feature value falls close to the category boundary) will have a stronger impact on choices than outlying flankers. This occurs because the context-dependent gain field is, on average over all trials, Gaussian with a mean of approximately zero; in other words, the tuning of neurons near the category boundary is on average sharper. When we applied the same analysis to human data, we observed exactly the same pattern, with inlying flankers exerting a more distracting influence on choices **(Fig. 4a)**. Applying the same regression model to predict RTs, the plots of 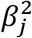 for both humans and the model show the characteristic repulsion effect in choices, that is owing to the greater contextual modulation of neural tuning near the category boundary **(Fig. 4b)**. Together, these analyses show that the human data resemble the counterintuitive model predictions over 4 different experimental datasets.

**Figure 4.**
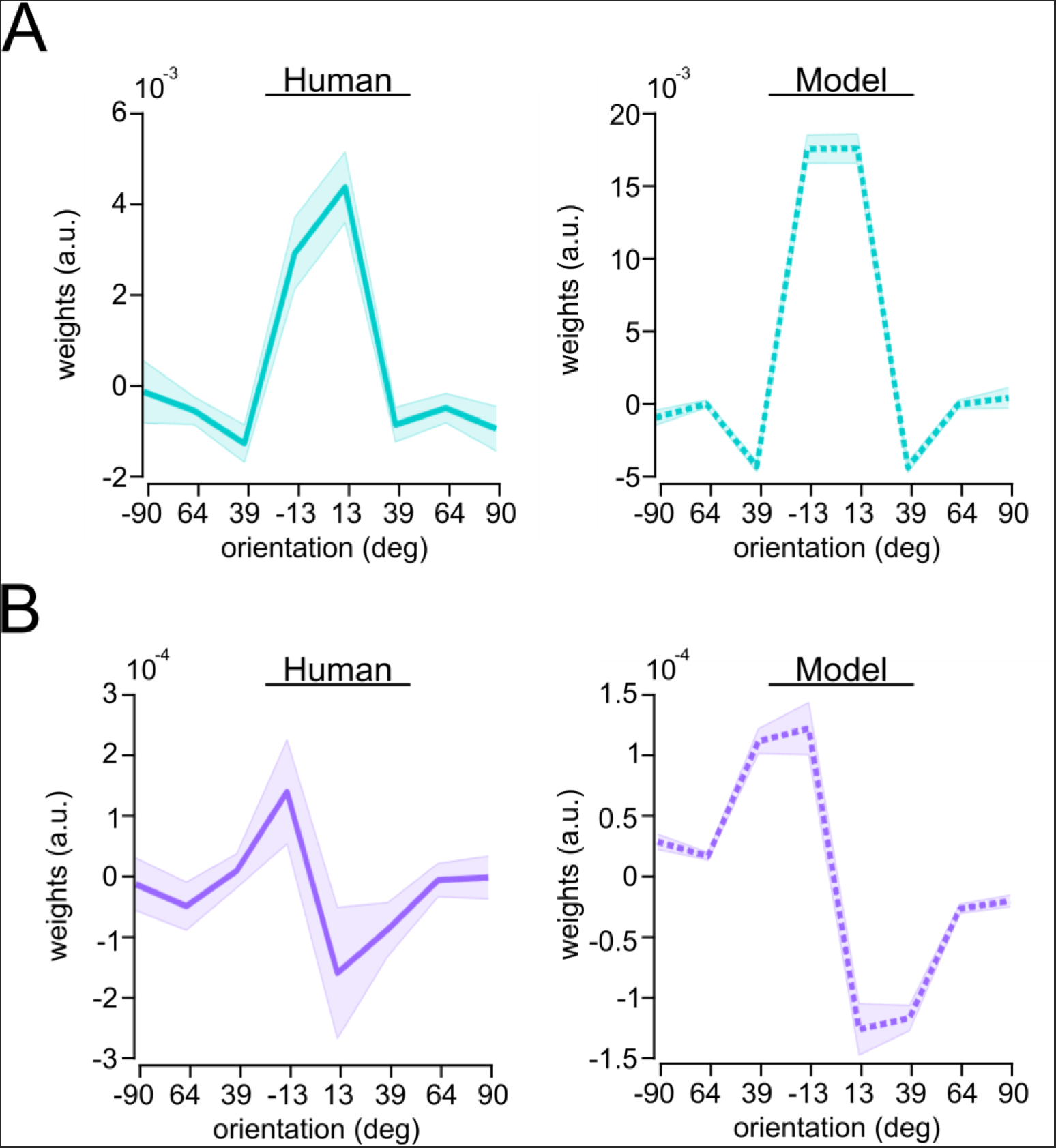
Choice bias driven by the flankers. Flanker tilts sorted into 8 bins from most CCW to most CW and used for predict choice (panel **A**; using probit regression) and response times (panel **B**) for both humans (left panels) and the best-fitting model (right panels). The x-axis shows the centre of each tilt bin, and the y-axis shows the resulting coefficients. The inverted-U-shape pattern in Panel A reveals that the inlying flankers have a stronger impact on choice than outliers for both humans and model. The response time data reveal the repulsive nature of this effect (Panel B solid purple line; note similarities with **Fig. 1a**). Higher absolute magnitude of beta weights implies that orientations that are close to the category boundary are more predictive of RT, similar to what was shown on choice in panel A. Shaded areas are the standard error of mean.

### Functional brain imaging

Established theories propose that a brain network that prominently includes the dorsal anterior cingulate cortex (dACC) is involved in the recruitment of control processes that allow the brain to overcome distraction. Across a range of paradigms including the Eriksen Flanker task, the dACC responds with higher-amplitude BOLD signals on incompatible than compatible trials^35,36^, and this effect is accentuated when the previous trial was compatible (‘conflict adaptation’), as if the dACC is monitoring for conflict and signalling its onset^37^. However, the dACC is also implicated in decision processes more generally. For example, it signals the level of noise that corrupts an imperative stimulus during perceptual discrimination^38^, and its proximity to a choice point or category boundary^39^; it responds to the relative economic value of an unchosen to a chosen option^40-42^, to the value of switching to a new task or context^43,44^, and to the update signals that occur as decision values change^45^. The search for a unifying theory of the dACC has been one of the most challenging and controversial themes in cognitive neuroscience over recent years^28,46-48^.

To assess the role of the dACC and interconnected regions in adaptive gain control, we conducted a new experiment in which *X^i^*, *mX^j^* and *sX^j^* varied parametrically from trial to trial, rather than in a conditionwise fashion. A new cohort of humans (**Exp.3**; n = 20) performed this task whilst we acquired BOLD signals from across the brain using functional magnetic resonance imaging (fMRI). Behavioural results of this experiment replicated those from **Exp.2** (**Fig. S5**), and so we focussed on neural analysis to test whether brain signals indexed decision information in a way that was predicted by the adaptive gain model. We began by confirming previous reports that the dACC responds more vigorously when a target feature lies closer to a category boundary, i.e. in our experiment, when the target orientation is closer to vertical^39^. We first regressed |*X^i^*| (i.e. proximity of the target to the category boundary) alone against BOLD signals occurring at the time of choice across the entire brain (GLM1). Consistent with previous observations, we observed a negative effect of |*X^i^*| in the dACC (peak: 2, 8, 54, t_19_ = 9.66, p_fdr_ < 0.001), as well as the anterior insula (AIC; peak: 34,34,2, t_19_ = 10.11, p_fdr_ < 0.001) and superior parietal lobe (SPL; peak: 18, -68, 58, 119 = 8.09, p_fdr_ < 0.001 see Fig. 5a). Extracting regions of interest from these areas in a leave-one-out fashion across participants (see Methods), we then plotted how the BOLD signal varied in quartiles of both *X^i^* and *Xm^j^* (GLM2) and compared these signals to the predictions of (i) the adaptive gain model, (ii) an equivalent model with no adaptive gain, i.e. where all simulated cells had equivalent tuning width, and (iii) a model in which BOLD signals were driven by conflict alone **(Fig. 5b)**. We found that the pattern of BOLD signals in all 3 regions closely resembled that predicted by the adaptive gain model, but not the other models **(Fig. 5c)**. Specifically, although BOLD responses were elevated when the *X^i^* was close to zero (dACC: F_1,19_ = 52.37, p < 0.001; AIC: F_1,19_ = 53.4, p < 0.001; SPL: F_1,19_ = 48.94, p < 0.001), this effect was exaggerated on those trials where *mX^j^* was far from zero but of *compatible* sign (i.e. greater BOLD response in dACC, AIC and SPL on congruent relative to incongruent trials; dACC: t_19_ = 3.03, p = 0.0069; AIC: t_19_ = 2.82, p = 0.011; SPL: t_19_ = 2.28, p = 0.034). No such modulation was observed when *X^i^* was far from zero, as predicted by the adaptive gain model.

**Figure 5.**
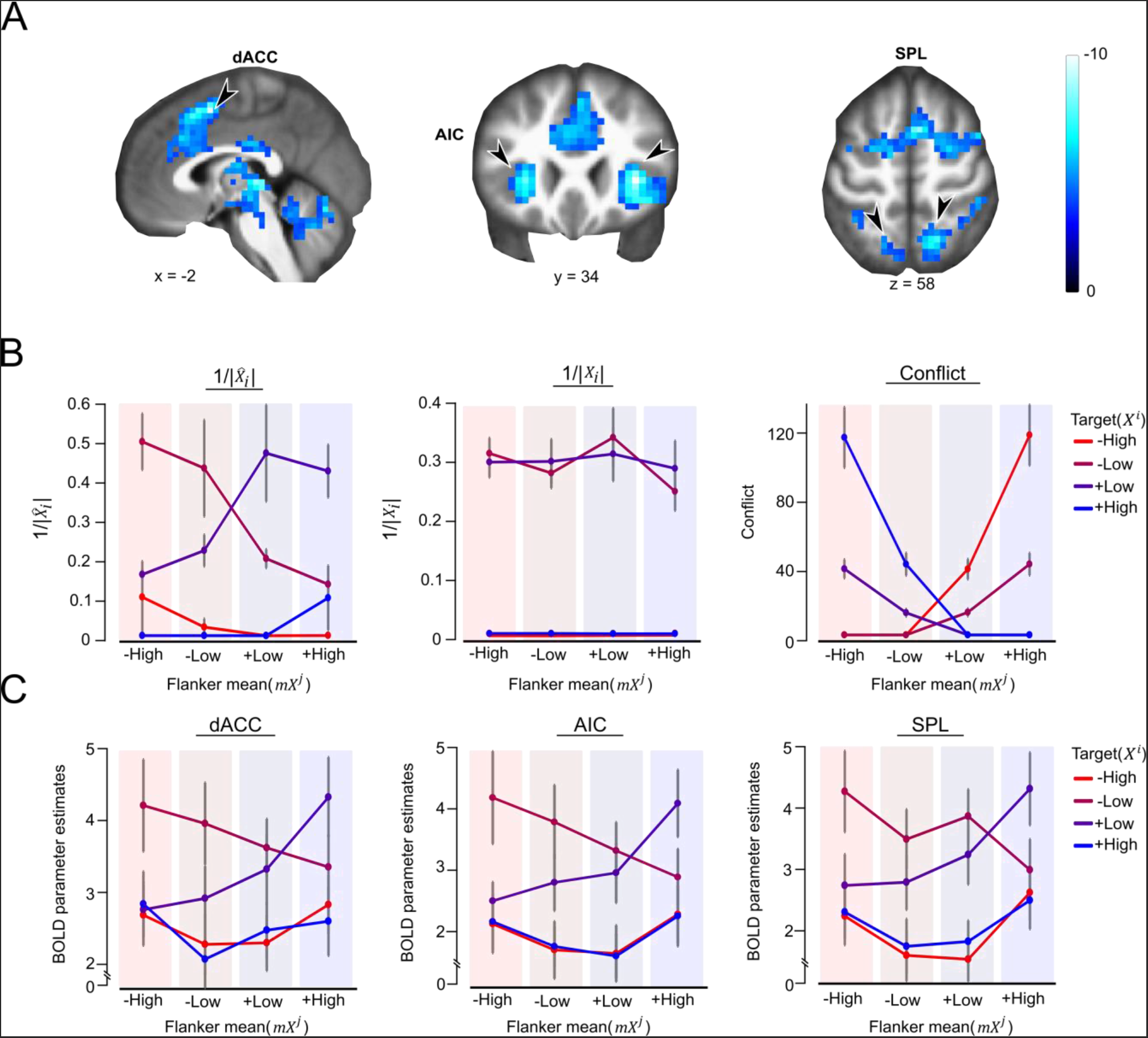
Effect of target orientation on BOLD signal. **(A)** Brain areas correlating negatively with the absolute target decision variable, rendered onto a template brain in sagittal (left panel) and axial (middle and right) slices. Images were generated with an uncorrected threshold of p < 0.0001. **(B)** Parametric modulators *X^i^* and *mX^j^* were each being split into 4 quartiles (lines for *X^i^*, x-axis or shaded area for *mX^j^*). Mean predictions (±SEM) from different models: Reciprocal gain-modulated decision variable 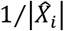 (left panel). Reciprocal no-gain modulated target decision variable 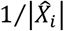 (middle panel). Estimated conflict was plotted for from each quartile bins (right panel). Coloured lines correspond to 4 levels of |*X^i^*|. Shaded coloured background (groupings on x-axis) corresponds to 4 levels of *mX^j^* **(C)** Mean BOLD (±SEM) signal beta values for each level of |*X^i^*| and *mX^j^* from the quartile bins. Left: dACC, Middle: AIC, and Right: SPL ROIs.

This suggests that a gain-modulated decision variable, rather than a conflict signal per se, is driving the dACC response. However, to quantify and compare the predictions of different models, we used Bayesian neural model comparison^49^. We fit the adaptive gain model and the rival conflict models on the trial RTs. Model estimates |*X^i^*| and conflict computed in different ways (see Methods) from the best fitting parameters are then used to estimate BOLD signals. We computed, within the dACC, AIC and SPL, the posterior probability of the adaptive gain model conditioned on the BOLD signal using Random effects Bayesian model selection from the VBA toolbox^50^, and compared the resulting estimates to those obtained for rival models. Both the exceedance probabilities and the expected frequencies strongly favoured the adaptive gain model over a decision model with no gain modulation, as well as over a family of conflict models (exceedance probabilities for the adaptive gain model in dACC: 0.992; AIC: 0.994; SPL: 0.996; expected frequencies for the adaptive gain model compared to chance level in dACC: t_19_ = 4.11, p < 0.001; AIC: t_19_ = 4.11, p < 0.001; SPL: t_19_ = 4.76, p < 0.001; **Fig. 6a**). In other words, the dACC, along with AIC and SPL, code for a decision signal modulated in precisely the fashion predicted by the adaptive gain model.

Finally, we addressed a concern that dACC is simply exhibiting a BOLD signal that correlates with the response production time on each trial^51^. Disentangling these factors is challenging, because (as described above) the model does an excellent job of predicting RTs. Nevertheless, when we included both model output 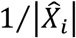 and RTs as competitive predictors in the model (GLM4), we still recovered a significant activation in the dACC (peak: 2, 20, 50, t_19_ = 5.28, p_fdr_ = 0.023), and AIC (left peak: -30, 20, 6, t_19_ = 6.09, p_fdr_ < 0.001; right peak: 34, 24, -6, t_19_ = 5.94, p_fdr_ < 0.001). In other words, the dACC BOLD signal correlates better with the demand predicted by the adaptive gain model than it does with time taken to produce a response on each trial.

How does our model explain previously-reported findings, such as the observation that the dACC responds to conflict^52^, or to the relative value of an unchosen vs. a chosen option during economic choice^53^? We have already shown that in the simple version of the Flanker task (c.f. **Exp.1a** where *sX^j^* = 0), the model predicts a larger output 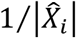 on incompatible relative to compatible trials. The model is thus in clear accord with a large literature indicating that dACC BOLD increases when target and distracters are incongruent in a simple version of the Flanker task^54^. We note that as described here, the adaptive gain model computes decision values independently on each successive trial, and thus in its current form would not predict conflict adaptation in the dACC. However, one could reasonably assume that adaptive effects may spill over from one trial to the next, i.e. that neural tuning width will be partly modulated by the previous trial. Under this assumption, the adaptive gain model will successfully predict that responses should be faster on two successive incongruent or two successive congruent trials^55^, just as it successfully account for the observation that during categorisation of a multi-element array, response times are faster if the target array is preceded by a prime array with an equivalent level of feature variance^29^.

However, we also note another facet of our results: that BOLD signals in the dACC, AIC and SPL ROIs correlate negatively with |*X^i^*| but positively with |*mX^j^*|(GLM5; **Fig. 6b**; dACC: t_19_ = 2.27, p = 0.035, AIC: t_19_ = 3.03, p = 0.003, SPL: t_19_ = 4.56, p = 0.007; this effect was also significant at the whole-brain level in voxels within the AIC and SPL, but not dACC). If we consider the target to be a “chosen” option and the flankers as a competing, “unchosen” option, the ensemble of findings reported here is reminiscent of the well-described finding by which dACC signals scale positively with the decision value associated with an unchosen option (i.e. the flankers) and negatively with the value of a chosen option (i.e. the target). Building on this intuition, we tested more directly the coding of model-predicted value of a chosen and unchosen option in a further simulation in which decision values for two stimuli were drawn randomly and independently from two distributions, and model output was converted to a choice via a softmax function (see Methods). This allowed us to correlate model output (i.e. predicted BOLD) with the value of the chosen and unchosen option, revealing a negative correlation with the former and a positive correlation with the latter, as previously reported^56^ **(Fig. 6c)**. Our model thus unifies a number of disparate accounts that have emphasised a role for the dACC in tasks involving categorising perceptual stimuli, and choosing among economic prospects.

**Figure 6.**
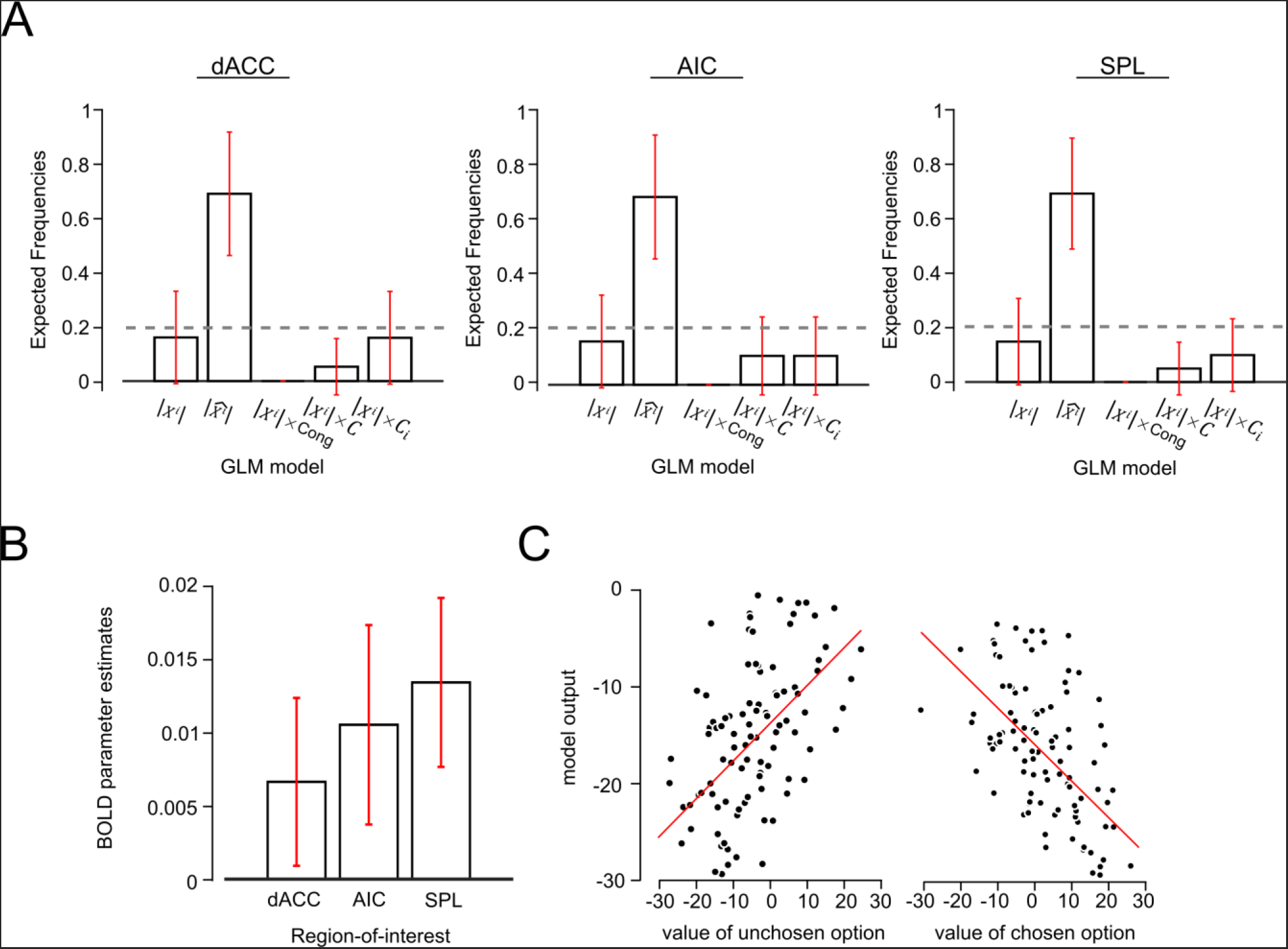
**(A)** Model comparisons across GLM models. Expected frequencies for each GLM and each ROI were computed using Random effects Bayesian model selection. Grey dotted line displayed the chance level **(B)** ROI analysis on dACC, AIC, and SPL on the absolute flanker mean decision variable |*mX^j^*|. Bar plots correspond to the mean (±SEM) beta estimates for this predictor from GLM5. **(C)** Simulated (negative) model output as a function of the value of chosen and unchosen option (see methods for details).

## Discussion

Good choices are based solely on information that is relevant to the choice at hand, and rational agents will successfully ignore distracting signals when making decisions^1^. However, across perceptual, cognitive and economic domains, human participants are observed to deviate from this rational principle. A range of different theories have been proposed to account for human susceptibility to distraction, but thus far, no single model has emerged that can account for phenomena as diverse as visual illusions, susceptibility to conflicting contextual features, or economic decoy effects. Here, we describe one such account. Our adaptive gain model has previously been shown to successfully account for diverse contextual influences in perceptual decision-making, including confirmatory biases in sequential sampling^23^ and priming by second-order summary statistics in perceptual categorisation^29^. Here, we show that not only can it account for classical effects of distraction across perceptual, cognitive and economic domains, but also that it successfully predicts a range of previously unreported, counterintuitive findings in a well-studied cognitive paradigm, the Eriksen flanker task^8^.

The effect of distraction is most often modelled under the assumption that irrelevant features are imperfectly filtered during decision-making, driving residual activation that corrupts decisions. When target and distracters prompt conflicting sensorimotor responses, the resulting competition slows response times and increases error rates^14^. A very successful neurocognitive theory proposes that dedicated processing systems have evolved in the primate medial prefrontal cortex that detect this conflict, and that are responsible for mobilising control mechanisms (associated with the lateral prefrontal cortex) to help mitigate the resulting costs^57^. Here, we propose a differing view: one that emphasises the benefits of consistent context rather than the costs of inconsistent context. In the adaptive gain model, contextual features offer guidance as to where to best allocate gain across feature space, ensuring that neurons that code for the most prevalent (or “expected”) features have the sharpest tuning and thus provide the most sensitive outputs. The adaptive gain model is thus motivated by the more general view that the nervous system has evolved to code for efficiently for sensory inputs, reducing redundancy by dynamically adjusting the tuning properties of decision-relevant neurons to maximise sensitivity to expected features^24,58^. In the flanker task, thus, the difference between compatible and incompatible trials arises at least in part because of a contextual facilitation mechanism at the decision level, akin to that described in sensory circuits^59^, rather than because of an active cost of response competition. This idea is not without precedent in theories of control. In fact, the notion that a flexible response to stimulus conflict is dependent on adaptive expectation mechanisms dates back to the original discovery of conflict adaptation by Gratton and colleagues more than 25 years ago^55^.

We take the opportunity to highlight two major features of our behavioural data that cannot be accounted for by standard accounts that emphasise the cost of conflict alone. Firstly, in **Exp.1a-b**, we found that a low-variance flanker array hastens response times on congruent trials, rather than prolonging response times on incongruent trials. This is consistent with an account that emphasises the benefit of consistent context rather than the cost of inconsistent context. Secondly, in **Exp.2a-b**, we observed that the longest response times were in fact observed on compatible trials, not incompatible trials. We replicated this finding across two different classes of visual feature: tilt and colour. According to our model, this cost occurred when the gain field dictated by the context repulsed the target subjectively closer to the category boundary, rendering choices more uncertain. Although such negative flanker effects have been reported with heavily masked stimuli, where they can be explained by differing timecourses of facilitatory versus inhibitory processes^60^, only rarely have such phenomena been reported for fully visible targets and distracters such as ours. Most interestingly, one such report occurred for a modified version of the flanker task where the targets were letters that were parametrically morphed between two possible identities, each corresponding to a possible flanker^61^. This report describes negative flanker effects when the target is most ambiguous, precisely parallelling our findings here for trials with small *X^i^* and large m*X^j^*, and a shift in the psychometric function that occurs with flanker identity in precisely the fashion predicted by our adaptive gain model^23^.

Our behavioural findings were echoed in the neural data recorded from dACC, where BOLD signals were higher when targets fell closer to the category boundary, but these signals were positively modulated (yet higher) when the distracters mean was congruent but further from the boundary. Without further assumptions, a model based on conflict alone cannot account for these findings. We do not wish to argue that stimulus or response conflict do not ever incur an additional cost to accuracy and response times, or that such a cost is unable to drive the dACC. Nevertheless, in the current study, we found that such an account was not required to explain our data, and that a model embodying this assumption fit our data more poorly.

Our findings present a challenge to extant theories, but we acknowledge that our model is currently incomplete. A large literature implicates the dACC in the mechanisms by which we update the value of actions in dynamically changing environments^45,62^. Our experiments were conducted in stationary settings, and we do not doubt that these regions may play additional roles (potentially also related to gain control) when slower learning about a changing context is required. We also note an important shortcoming in our findings: we were unable to identify differing roles for the dACC, AIC and SPL, which seem to act as one in our study. We think it is likely that our BOLD data are simply indexing the output of a decision process that involves modulation by distracting context, but are unable to make strong claims about the interim processes by which the computations proposed by the model occur. We suspect that exploring the role of adaptive gain control in dynamically-changing environments may shed more light on the differing contributions made by these regions, and we hope to pursue this question in future studies.

## Methods

For behavioural studies **Exp.1** and **2**, human participants were recruited via the online testing platform provided by Amazon Mechanical Turk (**Exp. 1a**: n = 37, 19 men, mean age: 31.8 years, SD = 8.09; **Exp.1b**: n = 36, 22 men, mean age: 31.12 years, SD = 9.57; **Exp. 2a**: n = 28, 14 men, mean age: 33.5 years, SD = 13.09; **Exp.2b**: n = 30, 16 men, mean age: 33.3 years, SD = 9.5). Mind here that the age information was collected by sets of age (e.g. 18-20, 21-30, 31-40 etc.) due to online testing, thus the reported age from Exp. 1a,1b,2a,2b was computed by taking the bin center of each age bin as the approximation of participants’ true age. For **Exp.3**, 20 healthy volunteers (8 men, mean age: 24.25 years, SD = 4.61) with normal or corrected-to-normal vision and no history of neurological disorders were recruited to participate. All participants gave informed consent to participate in the study and were compensated at a rate of $6 per hour for **Exp. 1** and **2** and €10 per hour for the fMRI scanning session. All experiments were approved and conducted in accordance with the Medical Sciences Inter-divisional Research Ethics Committee (MS IRDREC) ethical guidelines.

### Task details

#### Experiment 1a, 1b and 2a

Each trial began with a fixation dot 700ms, followed by the simultaneous presentation of target and flankers for 1000 ms. Target and flankers were full-contrast square-waved gratings with a 180° phase (5 cycles, and 75 pixels in diameter). The flankers are presented within a circular aperture of the target grating with 1.5 pixels separation with adjacent flankers. The orientation of each individual grating in the flanker surround was chosen form a Gaussian distribution with a particular mean (*mX^j^*) and standard deviation (*sX^j^*; see main text for precise values). To ensure that the sampled orientations matched the expected distribution with the given *mX^j^* and *sX^j^*, resampling of orientation values occurred until the mean and standard deviation of orientations fell within 1° tolerance of the desired *mX^j^* and *sX^j^*. Within the stimulus presentation duration, participants had to indicate whether the central grating was clockwise or counterclockwise to the vertical axis by pressing the right and left keyboard arrows respectively. They were instructed to ignore the flanking gratings. Visual feedback was given immediately after their response. Participants completed 6 blocks of 96 trials for **Exp. 1a**, 10 blocks of 96 trials for **Exp. 1b**, and 10 blocks of 108 trials for **Exp. 2a**).

#### Experiment 2b

While keeping all other aspects of the experiment (e.g. timing) the same, in **Exp.2b** we substituted the grating stimuli with coloured stimuli from a continuous colour space. Target and flankers were circles (100 pixels diameter per circle; 19 pixels separation with adjacent flanking circles) with hue ranging from blue to red. Participants were asked to indicate whether the central colour circle was more red (right arrow) or blue (Left arrow) while ignoring the surrounding coloured circles. Individual colour values (*c*) were defined by RGB values using expression: RGB = [*c*, 0, 1-*c*] with the category boundary being defined as *c* = 0.5 or RGB = [0.5, 0, 0.5]. The colour value for the central target stimulus and the flanker mean colour value were drawn from the set {0.35, 0.4, 0.45, 0.55, 0.6, 0.65}. To facilitate comparison between **Exp.2a** and **Exp.2b**, we recentered the category boundary value to 0, which means the central target colour values (*X^i^*) can be redescribed as ±0.05, 0.1, 0.15 with respect to a category boundary of zero. In this space individual flanker values were drawn from Gaussian distribution with means (*mX^j^*) of ±0.05, 0.1, 0.15 and standard deviations (*sX^j^*) of 0, 0.1, 0.2. Resampling of flanker feature values occurred until *mX^j^* and *sX^j^* fell within 0.01 from the desired values. Participants completed 10 blocks of 96 trials for this experiment.

#### Experiment 3 (fMRI study)

All stimuli were presented on a grey background. Participants first saw a fixation dot for 500ms, followed by the presentation of stimuli for 500ms, and were asked to respond within 1500 ms of stimulus onset. Stimuli were tilted gratings as for **Exp.1** and **Exp.2a**. At the end of response window, visual feedback in the form of a green or red dot indicated whether the response was correct or not. There was a jittered interval of between 2-6s (on average 4s) interposed between trials. Target tilt and flanker tilt statistics were varied independently on a trial-by-trial basis. On each trial, tilts of the target grating and flanker mean orientation were drawn independently from uniform distributions with a limit of -45° and +45°. Similarly, flanker standard deviation could be values between 0 to 30° from a uniform distribution. Resampling was used to ensure that the number of congruent and incongruent trials was matched across the entire experiment. Participants completed 4 (n = 10) or 5 (n = 10) blocks of the task in the scanner, with 130 trials on each block.

#### fMRI acquisition and preprocessing

Images were acquired with 3-Tesla Siemens TrioTim with a 32-channel head coil using a standard echo-planar imaging sequence. Images were 64 × 64 ×32 volumes with voxel size 3.5×3.5×3.5 mm; acquired with a 2s repetition time and echo time of 30 ms. We acquired fMRI data in 4 or 5 runs with 425 volumes per run. All preprocessing and fMRI analyses were carried out using SPM12. Preprocessing of the imaging data includes realignment of function images, co-registration of anatomical scan to the mean functional image, followed by segmentation and spatial normalisation to the standard template brain of the Montreal Neurological Institute (MNI brain). Lastly function images were spatially smooth with a 6-mm full width half maximum Gaussian kernel. A 128-s temporal high-pass filter was applied to exclude low-frequency artefacts.

#### Design and behavioural analysis

For experiments 1 and 2, the design orthogonalised the manipulation of target feature value (*X^i^*), mean of flankers (*mX^j^*) and variability of flankers (*sX^j^*). We can further designate trials as “congruent” when *X^i^* has the same sign as *mX^j^* or “incongruent” when *X^i^* has the opposite sign as *mX^j^*. In Experiment 1, we thus have a 3 × 2 (flanker variability × congruency) within-participant factorial design. In experiment 2, we introduced three levels of |*mX^j^*| and three levels of |*sX^j^*|, resulting in 3 × 3 × 3 × 2 (|*X^i^*|× |*mX^j^*| × *sX^j^* × congruency) within-participant factorial design with 54 conditions. Full list of *X^i^*, *mX^j^*, and *sX^j^* levels were displayed in **Table S1**. For **Exp. 1** and **2**, ANOVAS with Greenhouse-Geisser correction for sphericity were carried out at group-level analyses. A threshold of *p* < 0.05 was adopted for all behavioural analyses. We only analysed RT from correct trials, and additionally excluded trials where RT was faster than the 1% percentile or slower than the 99% percentile of the RT distribution. We used the same exclusion criteria across experiments.

### Computational modelling

#### Adaptive gain model

The computations that describe the population coding version of the adaptive gain model are described in the main text. To fit model outputs to human response time data (i.e. on a common scale in ms), for each parameterisation we regressed inverse decision values against each individual participants’ response times:

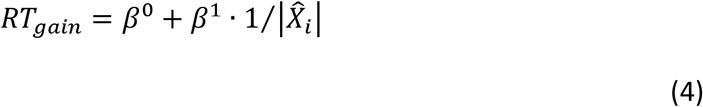

This calculation of RT is equivalent to modelling the data with ballistic (noiseless) diffusion process, with the two additional parameters *β*^0^ and *β*^1^ encoding the drift rate and non-decision time respectively (fixed across conditions). Searching, exhaustively across values of *σ^max^* and *ε* from equation (2), we identified the parameters that minimised mean squared error (MSE) between the human and model-predicted average RTs for each condition.

#### Conflict models

For the conflict model, we use a formulation described previously, whereby conflict *C* depends on the weighted product of competing inputs for the two actions *A^CW^* and *A^CCW^*

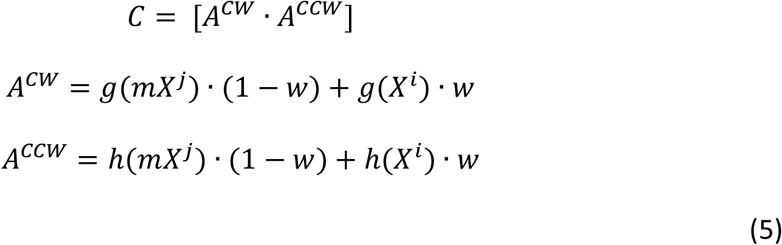

where *g(X)* and *h(X)* respectively denote positive and negative linear rectification functions. In conflict model 1, activation for the two actions *A^CW^* and *A^CCW^* is proportional to the tilt of the target and flanker mean. Alternatively, it can also be defined according to the tile of the target and each individual flanker (conflict model 2).

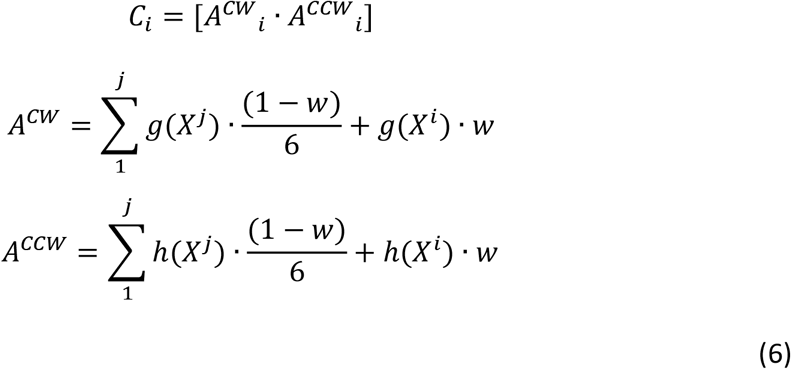

Where *j* is the number of flankers that are either congruent or incongruent. In other words, these models made different assumptions about conflict: that it was computed at the level of individual flankers (conflict model 2) or at the level of summary statistics (conflict model 1).

Finally, we compute RTs in a similar fashion as for the gain model:

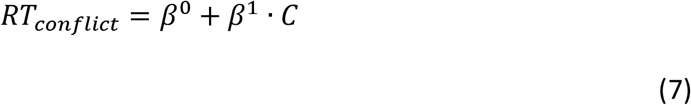

To reduce the risk of overfitting we used cross-fitting, estimating model parameters on half the trials and computing mean squared error (MSE) for the other half. These MSE were then fed into Bayesian Model Selection (BMS) to compute exceedance probabilities.

### Simulations of qualitative effects

When fit to human data, the adaptive gain model contained 2 free parameters: maximum tuning width (*σ^max^*) and sensory noise (*ε*). Simulations of the model aimed at qualitatively recreating effects from the past literature (e.g. for Fig.1, Fig. 6) assumed a fixed *σ^max^* (*σ^max^* = 10) and a fixed *ε* (*ε* = 5) unless noted otherwise.

#### Tilt illusion

In this simulation, we plot the difference between the true target angle (here, zero) and the gain modulated decision value 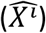 as a function of flanker mean decision value (*mX^j^* ∈ {−45, −44,…,+45°}) and flanker standard deviation (*sX^j^* ∈ {3,7,11,15}). For each variant of flanker mean decision values and flanker standard deviation, 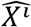 is computed using equation (1) and 2. We then plot 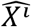 against levels of flanker mean decision value and flanker standard deviation in **Fig. 1a**.

#### Conflict effects

We computed a proxy of RT 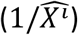 for the 3 conditions: ‘CO’: Congruent, where the target shares the same response association with the flankers; ‘SI’: Stimulus Incongruent, where the target is perceptually different to the flankers but the response associations of the two are still the same; ‘RI’: Response Incongruent, where the target has a different response association as the flankers **(Fig. 1b)**. In the simulation, flanker standard deviation *sX^j^* is set to be 0 in both ‘CO’ and ‘RI’ condition (i.e. we assumed flanker standard deviation was *ε*, or 5°). In ‘CO’, *X^i^* is equal to *mX^j^*(both are +45°). In ‘RI’, *mX^j^* has the opposite sign to the *X^i^*. Lastly, we simulated ‘SI’ condition by assuming *sX^j^* is higher than 0 i.e. *sX^j^* = 5; individual flankers are variable but *mX^j^* remained the same as *X^i^*. We assumed a higher maximum tuning width (*σ^max^* = 15) in this simulation.

#### Multi-alternative valued-based decision making task

In **Fig. 1c**, we simulated the difference between the model estimated decision values from the two targets (*X^i+^* = *20* & *X^1-^* = 10) as a function of a third distractors’ orientation (*X^j^* ∈ {−45, −44,…, +45°}). *sX^j^* was assumed to be 5 in this simulation. In **Fig. S1**, we computed the model output associated with the highest-valued choice-relevant alternative 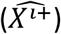 and the next best alternative 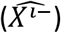, assuming that the mode of the gain field determined by the statistics lowest-valued (i.e. irrelevant) alternative *X^j^*. We then plotted a quantity proportional to choice probability, 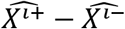, i.e. the relative difference in model output for the best and next best options.

#### Value of chosen vs. unchosen option

We simulated the model output as a function of the value of a theoretical chosen and unchosen option. On each trial, decision values for two stimuli (*X*^1^ and *X*^2^) were drawn independently from two zero-mean Gaussian distribution with a standard deviation of 10. On every trial, we allowed simultaneous evaluation of each stimulus in the context of the other, i.e. we passed each stimulus through the model as target with the alternative as distracter. We then assumed that participants chose according to the relative subjective (i.e. model output) value of the maximum and minimum resulting values, using a value of 5 for the slope of the choice function:

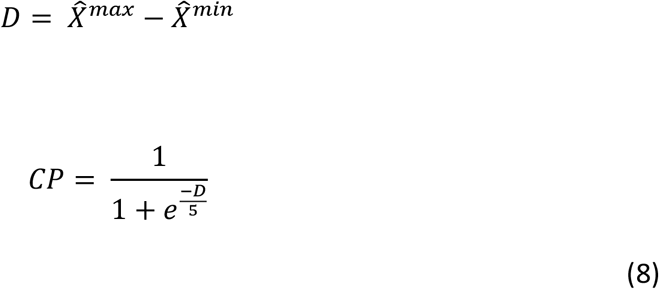

This allowed us to plot the relationship between *X* and model output *D* separately for the chosen and unchosen options.

#### Behavioural model comparison

We implemented Bayesian model selection at the group level to obtain posterior likelihoods and exceedance probabilities as described by Stephan and colleagues^63^.

### fMRI data analyses

#### Univariate Analyses

We analysed our data using Statistic Parametric Mapping (SPM12) with the general linear model (GLM) framework and in-house scripts running in Matlab. For all analyses, we ensured that sequential orthogonalisation of predictors in SPM was disabled. All GLMs also included regressors encoding the estimated movement parameters from preprocessing as a nuisance covariate. We modelled correct trials and trials that RT falls within 1 and 99 percentile distribution. All other trials were modelled separately as a nuisance regressor in all GLMs. We first constructed GLM1 with a single predictor encoding the parametric target decision values |*X^i^*|of the stimulus, time-locked to the onset of the stimulus. We identified voxels activated negatively by this regressor to define regions of interest in the dACC, AIC and SPL. Activations in these regions survived FDR correction for multiple comparisons at p < 0.05. To avoid double-dipping for later analyses, each region was identified in a leave-one-out fashion, with each participant in turn being omitted from a group-level analysis, which was used to define a Region-of-Interest (ROI) with a threshold of uncorrected p < 0.0001, from which beta values were extracted from the left out participant. For GLM2, we discretise each parametric modulator **(i)** target decision values |*X^i^*|and (ii) flanker mean feature value |*mX^j^*| into 4 quartiles bins and included a total of 4 x 4 = 16 regressors (corresponding to each quartile bins of feature values) in the GLM. BOLD betas from each subject are extracted using each ROI masks defined by the leave-one-out analysis. In GLM3, we entered the adaptive gain model estimates 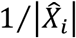 and binary regressor (congruency: +1 for congruent trials, -1 for incongruent trials) as competitive regressors. In GLM4, we replaced congruency with response time as a competing regressor to 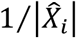. In GLM5, we included the following predictors as parametric modulators of the stimulus at the time of the decision **(i)** target decision values |*X^i^*|, (ii) flanker mean feature value |*mX^j^*| (iii) flanker precision, i.e. inverse flanker variability 1/*sX^j^*, (iv) absolute distance between target and flanker mean orientations |*X^i^* – *mX^j^*|, (v) the interaction of the distance between target and flanker mean with target decision values |*X^i^* – *mX^j^*| ⨯ |*X^i^*|, and lastly (vi) the interaction of the distance between target and flanker mean, target decision values and flanker variability (|*X^i^* – *mX^j^*| × |*mX^j^*| × 1/|*sX^j^*|). Full details of active voxels associated with these regressor were displayed in **Table S2-4**. Subsequent statistical tests on the betas extracted from each ROI were conducted using two-tailed one-sample t-test at a significance threshold of p < 0.05.

#### Neural Bayesian model comparison

We estimated the following GLMs using Bayesian statistics and conducted model comparisons to quantify and compare the predictions of different GLMs: **(i)** no-gain modulated decision value |*X^i^*| i.e. GLM1 described above, (ii) gain modulated decision value (|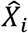|; GLM6), 3 GLMs that contained the no-gain modulated decision value and conflict defined in three ways (iii) GLM7: a binary congruency regressor (iv) GLM8: conflict computed using equation (4) (v) GLM9: conflict computed using equation (5). The fits of each GLM were assessed by Random Effects Bayesian Model Selection from the VBA toolbox^50^. This allows us to estimate the exceedance probabilities and expected frequencies for each model by feeding the log evidence for the aforementioned GLMs within each masks from the leave-one-out analysis. The best model is characterised with the highest exceedance probability and an expected frequency that is above chance (i.e. 1/number of models in comparison = 20%).

## Author Contributions

V.L., E.M. and C.S. conceived the study and designed the experiments; V.L., J.B. and S.H.C. collected data., V.L. and C.S. analysed the data, J.B. contributed custom scripts for fMRI data analysis. V.L. and C.S. wrote the manuscript, with helpful input from the other authors.

## Acknowledgements

We thank Tobias Egner for comments on the early version of the manuscript. This work was funded by European Research Council (ERC) Starter Grant (281628) to C.S., Croucher Foundation scholarship to V.L., Medical Research Council studentship to E.M. and the Wellcome Trust (0099741/Z/12/Z) to S.H.C. All funders had no role in the conceptualization, design, data collection, analysis, decision to publish, or preparation of the manuscript.

## Data Availability

All behavioural data are made available as Supplementary Information. Request for custom code and fMRI data should be directed to the corresponding author

## References

1 Luce R. D. Individual Choice Behavior: A Theoretical Analysis. (Wiley, 1959).

2 von Neumann, J. & Morgenstern, O. Theory of Games and Economic Behavior. (Princeton University Press, 1944).

3 Savage L. J. The Foundations of Statistics. (Wiley, 1954).

4 Wald A. & Wolfowitz J. Bayes Solutions of Sequential Decision Problems. Proc Natl Acad Sci U S A 35, 99–102 (1949).

5 Ma, W. J., Beck, J. M., Latham, P. E. & Pouget, A. Bayesian inference with probabilistic population codes. Nat Neurosci 9, 1432–1438, doi:nn1790[pii] 10.1038/nn1790 (2006).

6 Kording K. Decision theory: what ‘should’ the nervous system do? Science 318, 606–610, doi:10.1126/science.1142998 (2007).

7 Blakemore, C., Carpenter, R. H. & Georgeson, M. A. Lateral inhibition between orientation detectors in the human visual system. Nature 228, 37–39 (1970).

8 Eriksen B. A. & Eriksen C. W. Effects of noise letters upon identification of a target letter in a non-search task. Perception and Psychophysics 16, 143–149 (1974).

9 Kopp, B., Mattler, U. & Rist, F. Selective attention and response competition in schizophrenic patients. Psychiatry Res 53, 129–139 (1994).

10 Louie, K., Khaw, M. W. & Glimcher, P. W. Normalization is a general neural mechanism for context-dependent decision making. Proc Natl Acad Sci U S A, doi:1217854110[pii] 10.1073/pnas.1217854110 (2013).

11 Chau, B. K., Kolling, N., Hunt, L. T., Walton, M. E. & Rushworth, M. F. A neural mechanism underlying failure of optimal choice with multiple alternatives. Nat Neurosci 17, 463–470, doi:10.1038/nn.3649 (2014).

12 Tsetsos, K., Usher, M. & Chater, N. Preference reversal in multiattribute choice. Psychological review 117, 1275–1293, doi:10.1037/a0020580 (2010).

13 Tversky A. Elimination by aspects: A theory of choice. Psychological review 79, 281–299 (1972).

14 Botvinick, M. M., Braver, T. S., Barch, D. M., Carter, C. S. & Cohen, J. D. Conflict monitoring and cognitive control. Psychological review 108, 624–652 (2001).

15 Bundesen C. A computational theory of visual attention. Philos Trans R Soc Lond B Biol Sci 353, 1271–1281, doi:10.1098/rstb.1998.0282 (1998).

16 Heeger D. J. Normalization of cell responses in cat striate cortex. Vis Neurosci 9, 181–197, S0952523800009640 [pii] (1992).

17 Soltani, A., De Martino, B. & Camerer, C. A range-normalization model of context-dependent choice: a new model and evidence. PLoS Comput Biol 8, e1002607, doi:10.1371/journal.pcbi.1002607 PCOMPBIOL-D-12-00223[pii] (2012).

18 Schwartz, O., Sejnowski, T. J. & Dayan, P. Perceptual organization in the tilt illusion. J Vis 9, 19 11–20, doi:10.1167/9.4.19 (2009).

19 Dayan P. & Solomon J. A. Selective Bayes: attentional load and crowding. Vision Res 50, 2248–2260, doi:10.1016/j.visres.2010.04.014 (2010).

20 Logan G. D. The CODE theory of visual attention: an integration of space-based and object-based attention. Psychological review 103, 603–649 (1996).

21 Yu, A. J., Dayan, P. & Cohen, J. D. Dynamics of attentional selection under conflict: toward a rational Bayesian account. J Exp Psychol Hum Percept Perform 35, 700–717, doi:10.1037/a0013553 (2009).

22 Padoa-Schioppa C. Range-adapting representation of economic value in the orbitofrontal cortex. J Neurosci 29, 14004–14014, doi:29/44/14004 [pii] 10.1523/JNEUROSCI.3751-09.2009 (2009).

23 Cheadle S. et al. Adaptive gain control during human perceptual choice. Neuron 81, 1429–1441, doi:10.1016/j.neuron.2014.01.020 (2014).

24 Simoncelli E. P. & Olshausen B. A. Natural image statistics and neural representation. Annual review of neuroscience 24, 1193–1216, doi:10.1146/annurev.neuro.24.1.1193 (2001).

25 Rangel A. & Clithero J. A. Value normalization in decision making: theory and evidence. Current opinion in neurobiology 22, 970–981, doi:10.1016/j.conb.2012.07.011 (2012).

26 Kolling N. et al. Value, search, persistence and model updating in anterior cingulate cortex. Nat Neurosci 19, 1280–1285, doi:10.1038/nn.4382 (2016).

27 Shenhav, A., Cohen, J. D. & Botvinick, M. M. Dorsal anterior cingulate cortex and the value of control. Nat Neurosci 19, 1286–1291, doi:10.1038/nn.4384 (2016).

28 Heilbronner S. R. & Hayden B. Y. Dorsal Anterior Cingulate Cortex: A Bottom-Up View. Annual review of neuroscience 39, 149–170, doi:10.1146/annurev-neuro-070815-013952 (2016).

29 Michael, E., de Gardelle, V. & Summerfield, C. Priming by the variability of visual information. Proc Natl Acad Sci US A 111, 7873–7878, doi:10.1073/pnas.1308674111 (2014).

30 Solomon J. A. & Morgan M. J. Stochastic re-calibration: contextual effects on perceived tilt. Proc Biol Sci 273, 2681–2686, doi:10.1098/rspb.2006.3634 (2006).

31 Gibson J. J. Adaptation with negative aftereffect. Psychological review 44, 222–244 (1937).

32 Wu, T., Dufford, A. J., Mackie, M. A., Egan, L. J. & Fan, J. The Capacity of Cognitive Control Estimated from a Perceptual Decision Making Task. Scientific Reports 6 (2016).

33 de Gardelle, V. & Summerfield, C. Robust averaging during perceptual judgment. Proc Natl Acad Sci U S A 108, 13341–13346, doi:1104517108 [pii] 10.1073/pnas.1104517108 (2011).

34 Li V, Herce Castañón S, Solomon JA, Vandormael H, Summerfield C (2017) Robust averaging protects decisions from noise in neural computations. PLoS Comput Biol 13(8): e1005723. https://doi.org/10.1371/journal.pcbi.1005723

35 Van Veen V., & Carter, C.S. The timing of action-monitoring processes in the anterior cingulate cortex. Journal of cognitive neuroscience 14, 593–602 (2002).

36 Carter C. S. et al. Science. Science 280, 747–749 (1998).

37 Kerns J. G. et al. Anterior cingulate conflict monitoring and adjustments in control. Science 303, 1023–1026 (2004).

38 Ho, T. C., Brown, S. & Serences, J. T. Domain general mechanisms of perceptual decision making in human cortex. J Neurosci 29, 8675–8687, doi:29/27/8675 [pii] 10.1523/JNEUROSCI.5984-08.2009 (2009).

39 Grinband, J., Hirsch, J. & Ferrera, V. P. A neural representation of categorization uncertainty in the human brain. Neuron 49, 757–763, doi:S0896-6273(06)00089-4 [pii] 10.1016/j.neuron.2006.01.032 (2006).

40 Wunderlich, K., Rangel, A. & O’Doherty, J. P. Neural computations underlying action-based decision making in the human brain. Proceedings of the National Academy of Sciences 106, 17199–17204 (2009).

41 Tsetsos, K., Wyart, V., Shorkey, S. P. & Summerfield, C. Neural mechanisms of economic commitment in the human medial prefrontal cortex. eLife 3, e03701 (2014).

42 Juechems, K., Balaguer, J., Ruz, M. & Summerfield, C. Ventromedial Prefrontal Cortex Encodes a Latent Estimate of Cumulative Reward. Neuron 93, 705–714 (2017).

43 Kolling, N., Behrens, T. E., Mars, R. B. & Rushworth, M. F. Neural mechanisms of foraging. Science 336, 95–98 (2012).

44 Barber A. D. & Carter C. S. Cognitive control involved in overcoming prepotent response tendencies and switching between tasks. Cerebral Cortex 15, 899–912 (2004).

45 Rushworth, M. F., Noonan, M. P., Boorman, E. D., Walton, M. E. & Behrens, T. E. Frontalcortex and reward-guided learning and decision-making. Neuron 70, 1054–1069, doi:S0896-6273(11)00395-3 [pii] 10.1016/j.neuron.2011.05.014 (2011).

46 Shenhav, A., Botvinick, M. M. & Cohen, J. D. The expected value of control: an integrative theory of anterior cingulate cortex function. Neuron 79, 217–240 (2013).

47 Kolling, N., Behrens, T. E. J., Wittmann, M. K. & Rushworth, M. F. S. Multiple signals in anterior cingulate cortex. Current opinion in neurobiology 37, 36–43 (2016).

48 Shahnazian D. & Holroyd C. B. Distributed representations of action sequences in anterior cingulate cortex: A recurrent neural network approach. Psychonomic Bulletin & Review, 1–20 (2017).

49 Penny, W., Mattout, J. & Trujillo-Barreto, N. in Statistical Parametric Mapping: The analysis of functional brain images (eds K. Friston et al.) (Elsevier, 2006).

50 Daunizeau, J., Adam, V. & Rigoux, L. VBA: a probabilistic treatment of nonlinear models for neurobiological and behavioural data. PLoS computational biology 10, e1003441. (2014).

51 Grinband J. et al. The dorsal medial frontal cortex is sensitive to time on task, not response conflict or error likelihood. Neuroimage 57, 303–311, doi:S1053-8119(10)01610-1 [pii] 10.1016/j.neuroimage.2010.12.027 (2011).

52 Botvinick, M., Nystrom, L. E., Fissell, K., Carter, C. S. & Cohen, J. D. Conflict monitoring versus selection-for-action in anterior cingulate cortex. Nature 402, 179–181, doi:10.1038/46035 (1999).

53 Boorman, E. D., Rushworth, M. F. & Behrens, T. E. Ventromedial Prefrontal and Anterior Cingulate Cortex Adopt Choice and Default Reference Frames during Sequential Multi-Alternative Choice. J Neurosci 33, 2242–2253, doi:33/6/2242 [pii] 10.1523/JNEUROSCI.3022-12.2013 (2013).

54 Hazeltine, E., Poldrack, R. & Gabrieli, J. D. Neural activation during response competition. J Cogn Neurosci 12 Suppl 2, 118–129, doi:10.1162/089892900563984 (2000).

55 Gratton, G., Coles, M. G. & Donchin, E. Optimizing the use of information: strategic control of activation of responses. J Exp Psychol Gen 121, 480–506 (1992).

56 Boorman, E. D., Behrens, T. E., Woolrich, M. W. & Rushworth, M. F. How green is the grass on the other side? Frontopolar cortex and the evidence in favor of alternative courses of action. Neuron 62, 733–743, doi:S0896-6273(09)00389-4 [pii] 10.1016/j.neuron.2009.05.014 (2009).

57 Egner T. & Hirsch J. Cognitive control mechanisms resolve conflict through cortical amplification of task-relevant information. Nat Neurosci 8, 1784–1790, doi:nn1594 [pii] 10.1038/nn1594 (2005).

58 Barlow H. in Sensory Communication (MIT Press, 1961).

59 Stemmler, M., Usher, M. & Niebur, E. Lateral interactions in primary visual cortex: a model bridging physiology and psychophysics. Science 269, 1877–1880 (1995).

60 Eimer M. Facilitatory and inhibitory effects of masked prime stimuli on motor activation and behavioural performance. Acta Psychol (Amst) 101, 293–313 (1999).

61 Rouder J. N. & King J. W. Flanker and negative flanker effects in letter identification. Percept Psychophys 65, 287–297 (2003).

62 Behrens, T. E., Woolrich, M. W., Walton, M. E. & Rushworth, M. F. Learning the value of information in an uncertain world. Nat Neurosci 10, 1214–1221, doi:nn1954 [pii] 10.1038/nn1954 (2007).

63 Stephan, K. E., Penny, W. D., Daunizeau, J., Moran, R. J. & Friston, K. J. Bayesian model selection for group studies. Neuroimage 46, 1004–1017, doi:10.1016/j.neuroimage.2009.03.025 (2009).

